# CSCN: Inference of Cell-Specific Causal Networks Using Single-Cell RNA-Seq Data

**DOI:** 10.1101/2025.10.09.681381

**Authors:** Menghan Wang, Junya Yang, Luyao Lyu, Jiaxing Chen

## Abstract

Understanding gene regulation is fundamental to deciphering the coordinated activity of genes within cells. Although single-cell RNA sequencing (scRNA-seq) enables gene expression profiling at cellular resolution, most gene network inference methods operate at the tissue or population level, thereby overlooking regulatory heterogeneity across individual cells. Recent approaches, such as Cell-Specific Network (CSN) and its extension c-CSN, attempt to construct gene networks at single-cell resolution, providing a more detailed view of the regulatory logic underlying individual cellular states. However, these methods remain limited by high false positive rates due to indirect associations and lack of directionality or causal interpretability. To address these issues, we propose the Cell-Specific Causal Network (CSCN) framework, which infers directed, cell-specific gene regulatory relationships by explicitly modeling causality. CSCN combines causal discovery techniques with efficient computation using kd-trees and bitmap indexing to perform conditional independence testing, yielding sparse and interpretable causal graphs for each cell that effectively suppress indirect and spurious associations. We demonstrate through simulations that CSCN significantly reduces false positives compared to existing methods. Furthermore, we evaluate the quality of the inferred causal networks via clustering on the Causal Katz Matrix (CKM), and CSCN outperforms CSN and c-CSN in distinguishing cellular states.

## Introduction

Understanding how genes interact to regulate cellular behavior is a fundamental goal in molecular biology. Gene-gene networks, such as gene regulatory networks (GRNs), provide a powerful framework to model the coordinated activities among genes. However, most existing methods construct these networks at the tissue or population level, implicitly assuming regulatory homogeneity across cells. Increasing evidence suggests that gene interaction patterns can vary significantly from cell to cell, influenced by intrinsic heterogeneity, developmental dynamics, and microenvironmental factors.

This realization has driven growing interest in constructing gene networks at single-cell resolution, aiming to capture the unique regulatory and functional landscape of each individual cell. Advances in single-cell RNA sequencing (scRNA-seq) (Juzenas et al., 2025; Bageritz and Raddi, 2019) now enable gene expression profiling at the single-cell level, facilitating computational reconstruction of gene-gene networks on a per-cell basis. Gene networks at single-cell resolution offer a more detailed view of gene regulation by capturing the underlying control logic that shapes cellular functions. They provide critical insights into how individual cells interpret environmental signals, maintain distinct identities, and undergo state transitions. Compared to raw expression profiles, such network-based representations tend to be more stable and biologically informative, making them especially valuable for downstream single-cell analyses and for achieving a deeper mechanistic understanding of cellular heterogeneity.

Building on this foundation, the Cell-Specific Network (CSN) framework (Dai et al.) was proposed to infer gene–gene association networks at single-cell resolution by capturing nonlinear dependencies and constructing the Network Degree Matrix (NDM) to replace raw expression data, thereby enhancing downstream analyses. However, since CSN relies solely on marginal independence tests, it cannot distinguish direct from indirect dependencies, often leading to false positives. For instance, two genes independently regulated by a shared transcription factor may appear connected despite lacking a direct interaction.

To distinguish direct and indirect dependencies, the conditional Cell-Specific Network (c-CSN) framework (Li et al.) extends CSN by incorporating conditional independence tests thus reducing false positives in per-cell networks. While c-CSN improves specificity, it remains limited in two factors. First, its single-gene conditioning strategy fails to resolve indirect links when multiple regulators jointly control gene pairs—for example, genes A and B co-regulated by factors X and Y may still appear spuriously associated. Second, c-CSN produces undirected networks, lacking both edge directionality and causal interpretability. For example, it cannot differentiate whether gene A regulates gene B, vice versa, or if their relationship is reciprocal; all scenarios are treated as equivalent, potentially obscuring true regulatory direction and reducing the biological validity of downstream analyses.

To address these limitations, we propose the Cell-Specific Causal Network (CSCN) framework, a novel approach that, for the first time, infers causal gene regulatory networks at single-cell resolution. CSCN constructs directed graphs to represent gene regulatory interactions by explicitly modeling causality rather than relying on mere associations. This allows CSCN to eliminate indirect and spurious links, producing sparse, interpretable, and biologically meaningful networks in single-cell resolution. The resulting causal perspective offers a more accurate and robust representation of gene regulatory dynamics, enhancing downstream interpretability. CSCN adapts the causal inference algorithm Peter Clark (Pearl, 2009) with gene-level conditional independence testing. To deal with the high computational demands of causal discovery, we incorporated kd-tree data structures and bitmap indexing to markedly accelerate the conditional independence testing for large-scale gene and cell data.

We first demonstrate through simulations that CSCN significantly reduces false positives compared to existing methods. To evaluate the quality of the inferred causal networks, we perform clustering based on the Causal Katz Matrix (CKM), a representation derived from each cell-specific causal graph. CKM quantifies gene regulatory importance using Katz centrality, capturing both direct and multi-step causal influences while preserving the original expression matrix’s dimensionality for compatibility with standard downstream analyses. Empirical results across multiple scRNA-seq datasets show that CKM-based clustering consistently outperforms conventional representations—including GEM, NDM, and CNDM—in terms of accuracy, robustness, and biological interpretability. Additionally, CSCN facilitates biomarker discovery by identifying causal genes linked directly to disease phenotypes, providing insights to disease mechanisms.

In summary, CSCN offers the first scalable framework for inferring directed, cell-specific gene regulatory networks, from which CKM is derived to support accurate, interpretable, and robust downstream analyses.

## Methods

### Construction of CSCN

We propose the CSCN framework to infer directed regulatory interactions from single-cell scRNA-seq data. The CSN method infers cell-specific gene networks by testing pairwise dependencies between genes, but it may overestimate network connectivity due to indirect effects. To address this issue, CSCN refines the dependence testing between two genes, A and B, by conditioning on additional genes. Unlike c-CSN, which conditions only on a single selected gene, CSCN considers a broader set of candidate genes, up to full conditioning on all other genes. This broader conditioning helps to better eliminate indirect dependencies in high-dimensional settings. The framework proceeds in three main steps. First, dependence testing is applied to identify the network skeleton for each cell. Second, to ensure that the resulting graph gives the causality direction, CSCN applies a causal orientation step based on the PC algorithm with Meek’s orientation rules. Finally, since using a large conditioning set greatly increases computational demands, CSCN incorporates algorithmic optimizations to improve efficiency and scalability.

### Skeleton construction

Given *m* genes (or gene modules) and *n* single cells, CSCN produces *n* partially directed acyclic graphs (PDAGs) for causality network for every cells, denoted by {*G*^(1)^, …, *G*^(*n*)^}. First, CSCN incorporates a preprocessing step to reduce dimension and remove noise in scRNA-seq data(see Supplementary Section S1.1). Then, for each cell *k* ∈ {1, …, *n*}, when we consider the interaction between two genes *x, y*, we define their conditioning gene set as Z = {*z*_1_, …, *z*_*r*_}. The conditioning gene set contains the candidate gene *z* that may have *x*− *> z* and *z*− *> y* transfer intereaction and influence the inference of the indirect edge between *x*− *> y*. Then we define the localized neighborhoods for each gene in the expression space. For cell *k*, the neighborhood of a gene is an interval of predefined size centered around the expression value of that gene in cell *k*. Using these neighborhoods, we count neighboring cells *k*^′^ that satisfy different co-occurrence patterns: 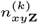 cells that simultaneously fall into the local neighborhood of *x,y* and Z, 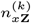 cells that fall into the local neighborhood of *x* and Z, 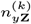 cells that fall into the local neighborhood of *y* and Z, 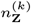 cells that fall into the neighborhood of Z. These counts are used to construct a localized deviation statistic, which measures the strength of the conditional association between x and y given Z:

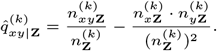

We then estimate the standard normal distribution of the count and test the p-value that a direct association exist between *x* and *y*. Under the null hypothesis *x* ⊥ *y* | Z, meaning there is no direct causal relationship between x and y after conditioning on Z, the normalized statistic 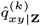 approximately follows a standard normal distribution. Edges are retained if the test statistic exceeds a predefined threshold: 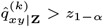, where *z*_1−*α*_ is the (1 − *α*) quantile of 𝒩 (0, 1). By iterating over conditioning sets of increasing size, CSCN progressively removes indirect connections, resulting in a pruned undirected network skeleton for each cell k. This refined skeleton forms the basis for the subsequent causal orientation step, ultimately producing a cell-specific PDAG.

### Orientation phase

Given the undirected skeleton, CSCN proceeds to orient edges by adapting the PC algorithm with Meek’s orientation rules. The orientation process consists of two main steps. First, v-structures are identified as follows. For every unclosed triple (*x* − *z* − *y*) with *x* and *y* nonadjacent, if *z* ∉ Sep(*x, y*), the triple is oriented as a collider *x* → *z* ← *y. Sep*(*x, y*) denotes the set of variables that rendered *x* and *y* conditionally independent during skeleton construction. This step ensures that minimal conditional dependencies detected during skeleton construction are faithfully encoded as converging causal arrows.After v-structures are identified, additional edge directions are inferred by iteratively applying Meek’s orientation rules under the constraint of acyclicity (Meek, 2013). In particular: (R1) if *a* → *b* and *b* − *c* with *a* and *c* nonadjacent, then orient *b* − *c* as *b* → *c* to avoid introducing fake colliders; (R2) if *a* − *c* → *b* and *a* − *b* is undirected, then orient *a* − *b* as *a* → *b* to maintain acyclicity; (R3) if two nonadjacent vertices *c* and *d* are both oriented as *c* → *b* and *d* → *b*, while *a* − *b, a* − *c*, and *a* − *d* are undirected, then *a* − *b* is oriented as *a* → *b* to preserve consistency. These rules are applied repeatedly until no further orientations can be made, yielding a maximally oriented graph for each cell. Edges that remain undirected after this process correspond to relationships whose direction cannot be uniquely determined from observational data alone. The resulting graph is a partially directed acyclic graph *G*^(*k*)^ per cell, capturing both the reliably inferred causal arrows and the undirected edges whose orientation cannot be resolved only on given data.

### Computational acceleration

Different with c-CSN that restrict Z to one conditional gene to reduce computational burden (Li et al.), CSCN explores larger conditioning sets, up to full conditioning on all other genes. This allows CSCN to more effectively eliminate indirect associations, but it also greatly increases computational demands. Naïve exhaustive conditional testing across *m* genes and *n* cells requires on the order of *O*(*m*^3^2^*m*^*n*) operations, as all possible conditioning sets must be enumerated. Such complexity is computationally prohibitive for large-scale single-cell datasets. To address this challenge, CSCN introduces several algorithmic optimizations to accelerate localized neighborhood queries and conditional probability estimation: First, gene expression profiles are indexed using a KD-tree, enabling neighborhood range queries to be performed in expected sublinear time *O*(*n*^1−1*/m*^)per query, rather than the naive linear *O*(*n*) cost. This significantly reduces the computational load in high-dimensional space. Second, conditional neighborhoods are cached using compact bitmap representations, allowing k-way set intersections to be executed via bit-parallel machine-word operations. This provides up to a 64 times speedup in practice when estimating conditional probabilities. Together, these optimizations (see Supplementary Section S1.2) yield order-of-magnitude performance improvements, making full conditioning across all genes computationally feasible. This enables CSCN to construct cell-specific causal networks while accounting for the complete set of genes, thereby improving the accuracy of inferred causal structures.

### Causal Katz matrix from CSCN

To quantify the causal influence of each gene, capturing both direct effects and indirect effects propagated through downstream regulatory pathways, we compute the Katz centrality for each cell’s CSCN. Katz centrality is a network measure that extends simple connectivity by accounting for all possible directed paths, while exponentially damping the contribution of longer paths. This makes it well-suited for modeling how perturbations to a gene may propagate through the regulatory network. For each cell k, the Katz centrality vector is defined as:

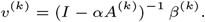

where *A*^(*k*)^ ∈ ℝ^*m*×*m*^ is the directed adjacency matrix of the inferred CSCN for cell *β*^(*k*)^ ∈ ℝ^*m*^ is a bias vector derived from the gene expression profile of that cell. The parameter *α* = 0.05 is a damping factor chosen to satisfy *α <* 1*/λ*_max_(*A*^(*k*)^), ensuring convergence of the inverse and controlling how strongly indirect effects are weighted relative to direct effects.

By concatenating the Katz centrality vectors across all cells, we construct the Causal Katz Matrix (CKM):

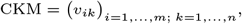

where each entry *v*_*ik*_ represents the causal propagation capacity of gene i in cell k. The CKM thus provides a cell-by-gene representation of causal influence, which can be directly integrated into downstream single-cell analyses such as clustering, trajectory inference, or identification of key regulatory drivers.

### Evaluation by clustering

We evaluated the clustering performance of CSCN using nine publicly available single-cell RNA-seq (scRNA-seq) datasets (Haber et al., 2017; Tasic et al., 2016; Camp et al., 2015; Close et al., 2017a; Leary et al., 2022; Segerstolpe et al., 2016; Fujimoto et al., 2020; Ranzoni et al., 2021).

Clustering was performed using four different types of input features: GEM (Gene Expression Matrix): the raw gene expression profiles. NDM (Network Degree Matrix): computed from the Cell-Specific Network (CSN). CNDM (Conditional Network Degree Matrix): derived from the conditional Cell-Specific Network (c-CSN). CKM (Card Katz Matrix), proposed in this study, represents each cell based on its inferred causal graph.

Two widely used unsupervised clustering algorithms, K-means and K-medoids, were applied to each feature type. The number of clusters was set to match the known number of cell types in each dataset. Clustering accuracy was measured by computing the Adjusted Rand Index (ARI). In addition, we compared runtime performance to evaluate the computational efficiency of CSCN relative to CSN and c-CSN. For visualization, UMAP(McInnes et al., 2020) and t-SNE(family=Maaten and Hinton) were applied to the feature matrices for dimensionality reduction. The resulting embeddings were plotted to qualitatively compare cluster separability across feature representations.

### Biomarker Discover

We applied CSCN to integrate breast cancer single-cell data (Xu et al.) with causal effect estimation to identify genes that have causal influences on diseases (Fig. 1). Gene expression profiles were obtained from cancer and healthy control samples, and CSCNs were constructed for each cell. These networks were then merged into a global causal network by including any edge present in at least one cell’s CSCN, ensuring broad coverage of potential causal interactions across diverse cell states.

**Fig. 1.**
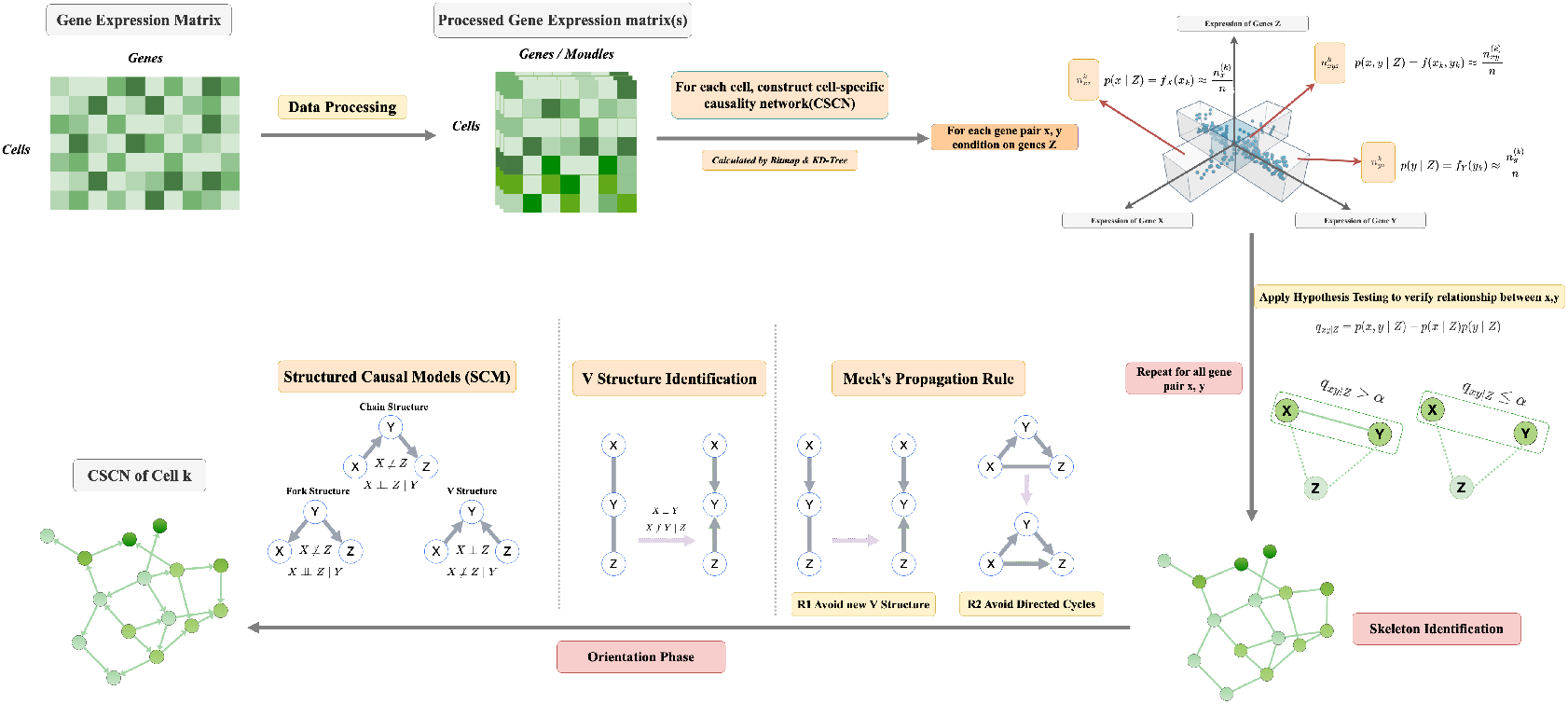
Workflow of CSCN.

To link genes to disease status, we added a virtual disease node, assigned a value of 1 for cancer cells and 0 for healthy cells. The goal is to estimate the Average Causal Effect (ACE) (Pearl, 2009; Pearl and Mackenzie, 2018) of each candidate gene *T* on disease outcome *Y*, while adjusting for confounders *C*, factors that influence both *T* and *Y* and can produce spurious associations. Using Pearl’s backdoor adjustment formula (Pearl, 2009; Pearl and Mackenzie, 2018), we can estimate the expected disease outcome under a hypothetical intervention on the candidate gene:

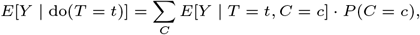

where *t* represents a specific expression level of *T*. Here, *E*[*Y* | *T* = *t, C* = *c*] is the conditional expectation of disease status given the gene expression and confounders, and *P* (*C* = *c*) is the probability distribution of the confounders. We approximate *E*[*Y* | *T* = *t, C* = *c*] nonparametrically using K-Nearest

Starting from single-cell expression matrices, preprocessing and dimensionality reduction (via NMF, WGCNA, or targeted subsets; see Supplementary Section S1.1) are applied. For each cell, conditional independence testing with full conditioning sets yields a pruned undirected skeleton. The orientation phase, implemented by the Peter–Clark algorithm, assigns causal directions to form a PDAG. CSCN employs KD-tree and bitmap-based accelerations (see Supplementary Section S1.2) to make full conditioning feasible, resulting in directed cell-specific causal networks.

### Neighbors (KNN)

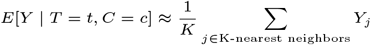

The ACE is then computed as the difference between interventional states:

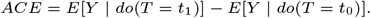

The terms *t*_0_ and *t*_1_ represent the interventional states of the gene’s expression, such as baseline versus elevated levels, allowing us to simulate the effect of altering the gene’s activity. A non-zero ACE indicates that perturbing the candidate gene’s expression directly alters the probability of the cell being in a disease state, even after adjusting for confounding factors. Conversely, an ACE near zero suggests that any association between the gene and disease status is likely non-causal or explained by other variables. Thus, genes with non-zero ACE values are considered causal biomarkers, as they play a direct and potentially actionable role in disease mechanisms rather than merely exhibiting correlative patterns.

## Result

### Simulation on confounding effects

We evaluated the proposed CSCN against the CSN and CCSN using a controlled simulation designed to test the impact of confounding regulation. In this simulation, four genes were modeled: two independent regulators (*Z*_1_, *Z*_2_ and two downstream genes (*X*, and *Y*). *Z*_1_ and *Z*_2_ were independently sampled from distinct normal distributions and jointly regulating *X* and *Y*. Importantly, there was no direct causal link between X and Y, meaning any inferred edge between them would represent a false positive caused by shared regulators.

We varied the regulatory strength of *Z*_1_ and *Z*_2_ from 0.1 to 2.0 and measured the false-positive rate (FPR) of each method (Fig. 2(A)). As the confounding influence increased, CSN’s FPR rose sharply, starting at 0.111 and reaching 0.878. CCSN performed better but still showed substantial confounding, with FPR increasing from 0.039 to 0.619. In contrast, CSCN maintained consistently low FPR, increasing only modestly from 0.027 to 0.148, even under the strongest confounding effects.

**Fig. 2.**
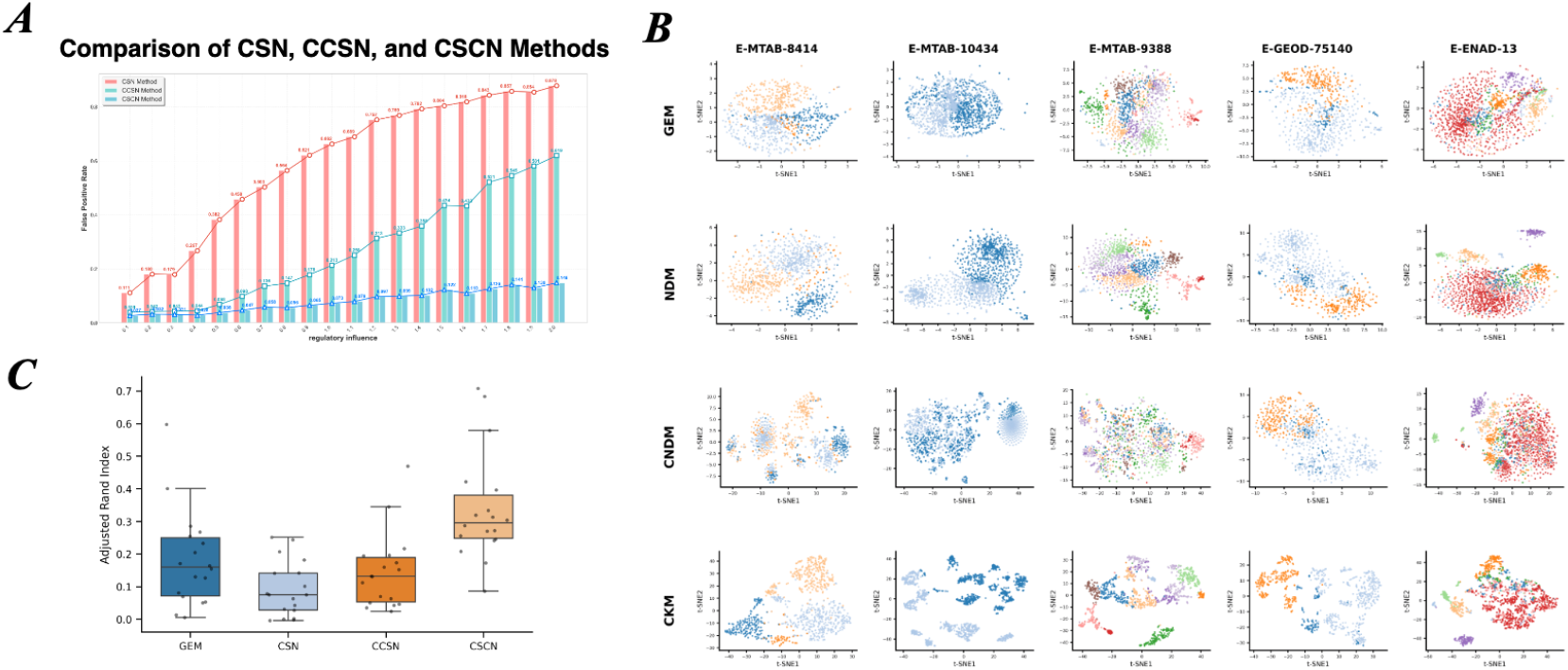
Clustering performance comparison across feature representations. (A) ARI scores using KMeans and KMedoids. (B) t-SNE visualizations of clustering structure. (C) ARI distributions across methods.

These results demonstrate that CSCN consistently achieves the lowest false-positive rate across all parameter levels, clearly outperforming both CSN and CCSN. This demonstrates CSCN’s superior specificity and robustness for accurate gene network inference in the presence of confounding regulation.

### Evaluation by clustering

To assess the quality of the inferred causal networks, we performed clustering analyses using the CKM representation across nine publicly available single-cell RNA-seq (scRNA-seq) datasets (Haber et al., 2017; Tasic et al., 2016; Camp et al., 2015; Close et al., 2017a; Leary et al., 2022; Segerstolpe et al., 2016; Fujimoto et al., 2020; Ranzoni et al., 2021).

CKM is derived from the CSCN by computing Katz centrality for each gene, which captures both direct regulatory effects and multi-step causal propagation. Using the K-means algorithm, CKM achieved the highest adjusted Rand index (ARI) on 6 out of 9 datasets (66.7%). With K-medoids, CKM ranks first on 7 out of 9 datasets (77.8%). Overall, across all 18 algorithm–dataset combinations, CKM achieved the top ARI in 13 cases (72.2%) (Supplementary Table 1).

To assess feature space separability, we visualized the embeddings using t-SNE (Fig. 2(B)). BCKM-based embeddings displayed clearer cluster boundaries, with reduced overlap between clusters and tighter within-cluster grouping. By contrast, representations from GEM, NDM, and CNDM showed greater inter-cluster mixing, consistent with their lower ARI scores(Fig. 2(C)). Aggregate ARI statistics are shown in boxplots (Fig. 2(C)). CKM not only achieved the highest median ARI but also exhibited the smallest interquartile range (IQR), indicating both better performance and greater stability across datasets.

### Temporal Dynamics on Network

Using dataset E-GEOD-93593, which profiles a 125-day differentiation protocol converting H1 human embryonic stem cells into various ventrally derived cell types (Close et al., 2017b), we identified three key gene modules involved in differentiation through WGCNA analysis (Langfelder and Horvath, 2008). We then examined their functional roles and regulatory rewiring across four developmental stages (Days 26, 54, 100, and 125) using CSCN.

At the global level, these modules showed clearly distinct functional enrichments (Fig. 3(A)). The turquoise gene module were enriched for cell migration, fatty acid metabolism, and extracellular matrix remodeling. Brown were primarily associated with neuronal development and synapse formation, whereas magenta were strongly linked to DNA replication and cell cycle regulation. These patterns indicate that each module represents a unique biological program during stem cell differentiation.

**Fig. 3.**
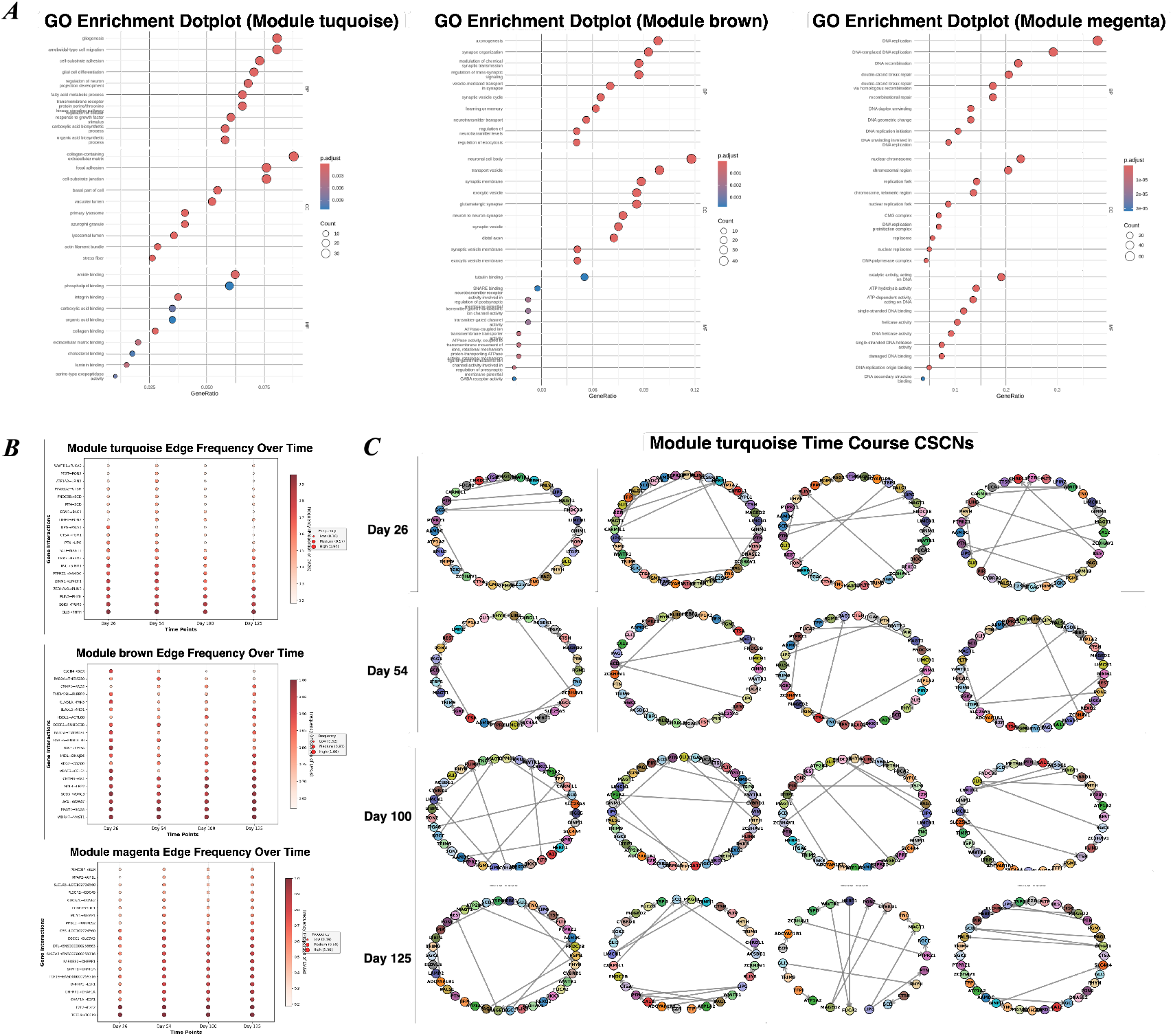
Network Rewiring Results. (A) GO enrichment dotplots showing key WGCNA modules. Dot size represents gene count; color indicates statistical significance (p-value). (B) SCP2046 Cell Network Entropy Results (C) Temporal dynamics of gene interaction frequencies in WGCNA modules. (D) Time-course cell-specific causal networks (CSCNs) for module turquoise.

Beyond static enrichment profiles, the functional activity of these modules shifted dynamically over time (Fig. 3(B)). Turquoise module activity intensified after Day 100, driven by interactions such as GLI3 → PHYH and SGK3 → TRIM9, reflecting late-stage structural remodeling. The brown module, linked to axon guidance and synaptic signaling, peaked at Days 100 and 125, with interactions like CLCN4 → DCX and PAK3 → CHGA becoming dominant. Meanwhile, the magenta module, associated with DNA replication and repair, surged at later stages through regulatory links such as TCF19 → TCF19 and E2F2 → E2F2. Together, these temporal dynamics suggest a sequential engagement of structural remodeling, neuronal maturation, and proliferative programs as differentiation progresses.

Our cell-specific causal network (CSCN) analysis uncovered that the gene regulatory networks (GRNs) of these modules are not uniform across cell types. For example, Fig. 3(B) illustrates how turquoise module interactions diverge between different cell populations, demonstrating distinct causal wiring despite shared module membership. This cell-type specificity underscores the advantage of our causal framework: it captures not only when modules are active, but also how their regulatory logic is rewired in a cell-specific manner. In summary, combining global enrichment (Fig. 3(A)), temporal dynamics Fig. 3(B), and cell-specific causal inference Fig. 3(C) provides a comprehensive view of network rewiring during stem cell differentiation into neurons.

In summary, CSCN integrates global enrichment, temporal dynamics, and cell-specific causal inference to provide a unified view of regulatory network dynamics.

### Biomarker Discovery Results

We applied the CSCN-ACE workflow to an initial list of differentially expressed genes identified by DESeq2, comparing breast tumor samples to healthy controls. By calculating the ACE, we filtered this list to isolate genes with a direct causal influence on disease state, resulting in a high-confidence set of 52 causal biomarkers. The overlap and distinction between the DESeq2 and CSCN-ACE results are shown in the Venn diagram (Fig. 4(E)). This process produced two gene groups for downstream analysis: the 52 causal biomarkers and 46 DESeq2-exclusive genes that were statistically significant but did not meet the causal criteria.

**Fig. 4.**
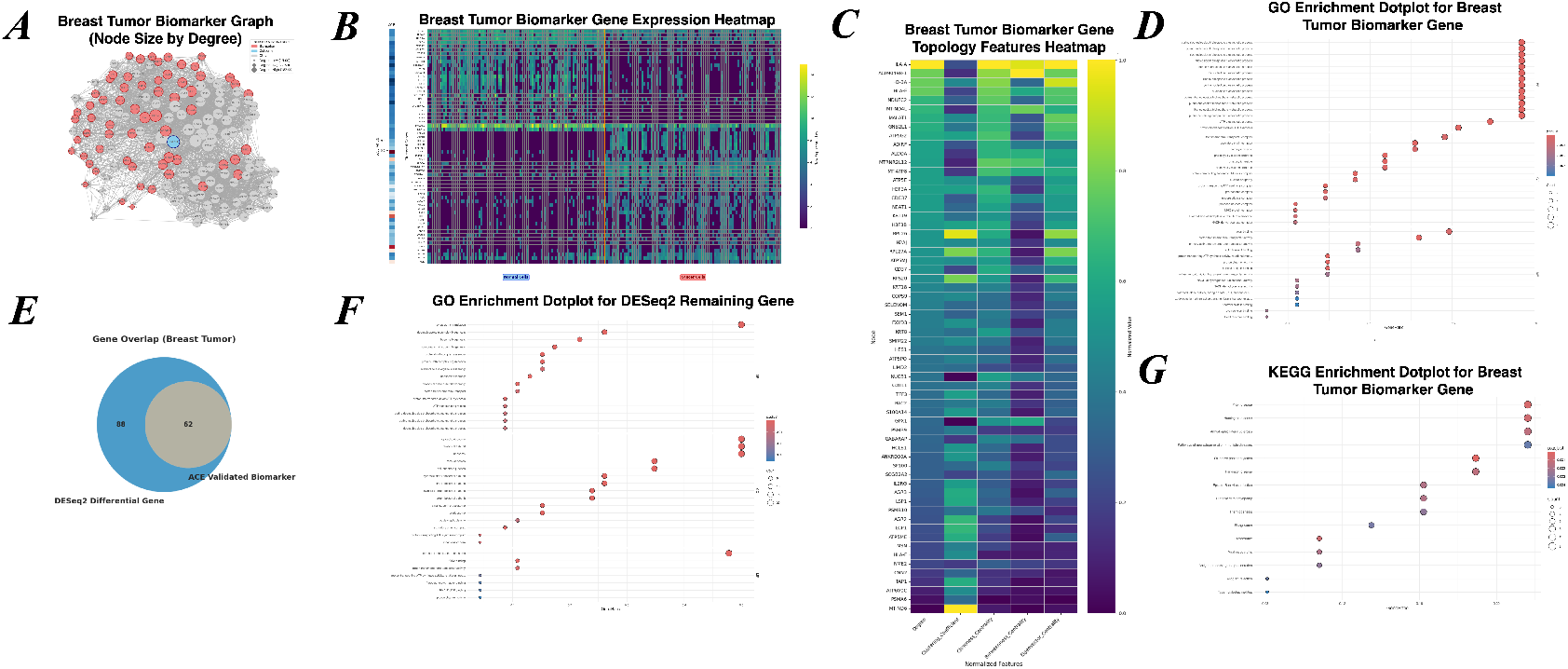
Biomarker Discovery Results. (A) Breast Tumor Biomarker Graph (Node Size by Degree). (B) Breast Tumor Biomarker Gene Expression Heatmap. (C) Breast Tumor Biomarker Gene Topology Features Heatmap. (D) GO Enrichment Dotplot for DESeq2 Remaining Gene. (E) Gene Overlap (Breast Tumor). (F) GO Enrichment Dotplot for Breast Tumor Biomarker Gene. (G) KEGG Enrichment Dotplot for Breast Tumor Biomarker Gene.

Functional enrichment analysis of the 52 causal biomarkers revealed a strong convergence on two fundamental hallmarks of cancer: metabolic reprogramming and immune evasion.

The KEGG pathway Oxidative phosphorylation (hsa00190, p.adjust = 9.82e-5) emerged as the most significantly enriched process (Fig. 4(G)). This finding was further supported by Gene Ontology (GO) analysis, which highlighted terms such as purine ribonucleoside triphosphate biosynthetic process (GO:0009206), closely linked to ATP synthesis (Fig. 4(D)). Concurrently, the KEGG pathway ‘Antigen processing and presentation’ (hsa04612) and the GO term ‘antigen processing and presentation of endogenous peptide antigen via MHC class I’ (GO:0019885) were also highly enriched, suggesting that these genes play a direct causal role in shaping tumor immune recognition.

Beyond functional enrichment, these biomarkers hold topological importance within the global causal network. They frequently appear as high-connectivity hubs (Fig. 4(A)) and exhibit elevated network feature scores (Fig. 4(C)), reinforcing their importance. Moreover, their expression profiles show consistent and marked dysregulation between tumor and control samples (Fig. 4(B)), reinforcing their biological relevance.

In contrast, the DESeq2-exclusive gene set was predominantly linked to basic protein synthesis. GO enrichment analysis for this group was dominated by ribosome-related processes, with cytoplasmic translation (GO:0002181, p.adjust = 9.10e-33) as the most significant term (Fig. 4(F)). This indicates that these genes primarily reflect the core machinery needed for rapid cell growth. Notably, strategic pathways, such as those involved in immune evasion, were entirely absent from this set. This sharp functional divergence highlights the distinction between traditional differential expression and causal inference. While DESeq2 captures the downstream effects of malignant proliferation, such as increased ribosome production, our causal method isolates the upstream regulatory drivers that actively shape the malignant phenotype.

## Conclusion

We introduced Cell-Specific Causal Networks (CSCN), a scalable framework for reconstructing directed causal gene regulatory relationships for each cell while reducing false positives. By translating causal topology into informative features such as the Causal Katz Matrix, CSCN improves clustering, and through measures like ACE, it enables causal biomarker discovery. Applications to scRNA-seq and spatial transcriptomics demonstrate its robustness and biological relevance. Despite remaining challenges, CSCN provides a foundation for interpretable causal analysis at single-cell resolution, offering insights into development, disease progression, and therapeutic targeting.7

## Acknowledgments

This work is supported by the National Natural Science Foundation of China (Grant No. 32200526) with match fund provided by BNBU Research Grant UICR0300014, and the Guangdong Provincial/Zhuhai Key Laboratory IRADS (2022B1212010006) and Guangdong Higher Education Upgrading Plan (2021-2025) with BNBU Research Grant UICR0400025-21, and BNBU Start Up Fund UICR0700039-22.

## References

J. Bageritz and G. Raddi. Single-cell rna sequencing with drop-seq. In Single Cell Methods: Sequencing and Proteomics, pages 73–85. Springer, 2019.

J. G. Camp, F. Badsha, M. Florio, S. Kanton, T. Gerber, M. Wilsch-Bräuninger, E. Lewitus, A. Sykes, W. Hevers, M. Lancaster, J. A. Knoblich, R. Lachmann, S. Pääbo, W. B. Huttner, and B. Treutlein. Human cerebral organoids recapitulate gene expression programs of fetal neocortex development. Proceedings of the National Academy of Sciences, 112(51):15672–15677, Dec. 2015. doi: 10.1073/pnas.1520760112.

J. L. Close, Z. Yao, B. P. Levi, J. A. Miller, T. E. Bakken, V. Menon, J. T. Ting, A. Wall, A.-R. Krostag, E. R. Thomsen, A. M. Nelson, J. K. Mich, R. D. Hodge, S. I. Shehata, I. A. Glass, S. Bort, N. V. Shapovalova, N. K. Ngo, J. S. Grimley, J. W. Phillips, C. L. Thompson, S. Ramanathan, and E. Lein. Single-cell profiling of an In Vitro model of human interneuron development reveals temporal dynamics of cell type production and maturation. Neuron, 93(5):1035–1048.e5, Mar. 2017a. ISSN 0896-6273. doi: 10.1016/j.neuron.2017.02.014.

J. L. Close, Z. Yao, B. P. Levi, J. A. Miller, T. E. Bakken, V. Menon, J. T. Ting, A. Wall, A.-R. Krostag, E. R. Thomsen, A. M. Nelson, J. K. Mich, R. D. Hodge, S. I. Shehata, I. A. Glass, S. Bort, N. V. Shapovalova, N. K. Ngo, J. S. Grimley, J. W. Phillips, C. L. Thompson, S. Ramanathan, and E. Lein. Single-Cell Profiling of an In Vitro Model of Human Interneuron Development Reveals Temporal Dynamics of Cell Type Production and Maturation. Neuron, 96(4):949, Nov. 2017b. ISSN 0896-6273. doi: 10.1016/j.neuron.2017.10.024.

H. Dai, L. Li, T. Zeng, and L. Chen. Cell-specific network constructed by single-cell RNA sequencing data. 47(11):e62. ISSN 1362-4962. doi: 10.1093/nar/gkz172.

p. d. u. family=Maaten, given=Laurens and G. Hinton. Visualizing data using t-SNE. 9(86):2579–2605. URL http://jmlr.org/papers/v9/vandermaaten08a.html.

N. Fujimoto, Y. He, M. D’Addio, C. Tacconi, M. Detmar, and L. C. Dieterich. Single-cell mapping reveals new markers and functions of lymphatic endothelial cells in lymph nodes. PLOS Biology, 18(4):e3000704. Apr. 2020. ISSN 1545-7885. doi: 10.1371/journal.pbio.3000704.

A. L. Haber, M. Biton, N. Rogel, R. H. Herbst, K. Shekhar, C. Smillie, G. Burgin, T. M. Delorey, M. R. Howitt, Y. Katz, I. Tirosh, S. Beyaz, D. Dionne, M. Zhang, R. Raychowdhury, W. S. Garrett, O. Rozenblatt-Rosen, H. N. Shi, O. Yilmaz, R. J. Xavier, and A. Regev. A single-cell survey of the small intestinal epithelium. Nature, 551(7680):333–339, Nov. 2017. ISSN 1476-4687. doi: 10.1038/nature24489.

S. Juzenas, K. Goda, V. Kiseliovas, J. Zvirblyte, A. Quintinal-Villalonga, J. Siurkus, J. Nainys, and L. Mazutis. indrops-2: a flexible, versatile and cost-efficient droplet microfluidic approach for high-throughput scrna-seq of fresh and preserved clinical samples. Nucleic Acids Research, 53 (2):gkae1312, 2025.

P. Langfelder and S. Horvath. WGCNA: An R package for weighted correlation network analysis. BMC Bioinformatics, 9(1):559, Dec. 2008. ISSN 1471-2105. doi: 10.1186/1471-2105-9-559.

N. Leary, S. Walser, Y. He, N. Cousin, P. Pereira, A. Gallo, V. Collado-Diaz, C. Halin, S. Garcia-Silva, H. Peinado, and L. C. Dieterich. Melanoma-derived extracellular vesicles mediate lymphatic remodelling and impair tumour immunity in draining lymph nodes. Journal of Extracellular Vesicles, 11(2):e12197, 2022. ISSN 2001-3078. doi: 10.1002/jev2.12197.

L. Li, H. Dai, Z. Fang, and L. Chen. C-CSN: Single-Cell RNA Sequencing Data Analysis by Conditional Cell-Specific Network. 19(2):319–329. ISSN 1672-0229, 2210-3244. doi: 10.1016/j.gpb.2020.05.005.

L. McInnes, J. Healy, and J. Melville. Umap: Uniform manifold approximation and projection for dimension reduction, 2020. URL https://arxiv.org/abs/1802.03426.

C. Meek. Causal inference and causal explanation with background knowledge. arXiv preprint arXiv:1302.4972, 2013.

J. Pearl. Causality. Cambridge university press, 2009.

J. Pearl and D. Mackenzie. The book of why: the new science of cause and effect. Basic books, 2018.

M. Ranzoni, A. Tangherloni, I. Berest, S. G. Riva, Myers, P. M. Strzelecka, J. Xu, E. Panada, I. Mohorianu, J. Zaugg, and A. Cvejic. Integrative single-cell rna-seq and atac-seq analysis of human developmental hematopoiesis. Cell Stem Cell, 28(3):472–487.e7, Mar. 2021. ISSN 1934-5909,1875-9777. doi: 10.1016/j.stem.2020.11.015.

Å. Segerstolpe, A. Palasantza, P. Eliasson, E. Andersson, Andréasson, X. Sun, S. Picelli, A. Sabirsh, M. Clausen, M. Bjursell, D. Smith, M. Kasper, C. Åmmälä, and R. Sandberg. Single-cell transcriptome profiling of human pancreatic islets in health and type2 diabetes. Cell Metabolism, 24(4):593–607, Oct. 2016. ISSN 1550-4131. doi: 10.1016/j.cmet.2016.08.020.

Tasic, V. Menon, T. N. Nguyen, T. K. Kim, T. Jarsky, Z. Yao, B. Levi, L. T. Gray, S. A. Sorensen, T. Dolbeare, D. Bertagnolli, J. Goldy, N. Shapovalova, S. Parry, C. Lee, K. Smith, A. Bernard, L. Madisen, S. M. Sunkin, M. Hawrylycz, C. Koch, and H. Zeng. Adult mouse cortical cell taxonomy revealed by single cell transcriptomics. Nature Neuroscience, 19(2):335–346, Feb. 2016. ISSN 1546-1726. doi: 10.1038/nn.4216.

L. Xu, S.-P. H. Huang, and I. Chan. Primary Breast Tumor Atlas. URL https://zenodo.org/records/10672250.

