## Supplemental for "CSCN: Inference of Cell-Specific Causal Networks Using Single-Cell RNA-Seq Data"

### S1. Methods

#### S1.1. Data Preprocessing

Depending on downstream goals and computational constraints, we apply one of three preprocessing strategies to focus and regularize causal inference:

##### *S1.1.1. Module-constrained gene-level inference via WGCNA.*

To recover full gene-level causal networks while avoiding wasted computation on biologically implausible edges, we first perform weighted gene co-expression network analysis (WGCNA) over all cells to partition genes into co-expression modules. Leveraging the sparsity of regulatory networks and the observation that regulatory interactions are enriched within co-expressed gene sets, we assume that causal edges do not exist between genes in different modules. Consequently, causal discovery is performed independently within each module, constructing gene-level subgraphs per cell. This modular constraint significantly reduces the search space by eliminating inter-module conditioning tests. To partially compensate for the representational loss induced by this assumption, we also infer a higher-level inter-module causal network by applying the CSCN procedure to module eigengenes, yielding a hierarchical (two-layer) causal representation in which module-level influences complement intra-module gene-level networks. The workflow is illustrated in Figure S1.

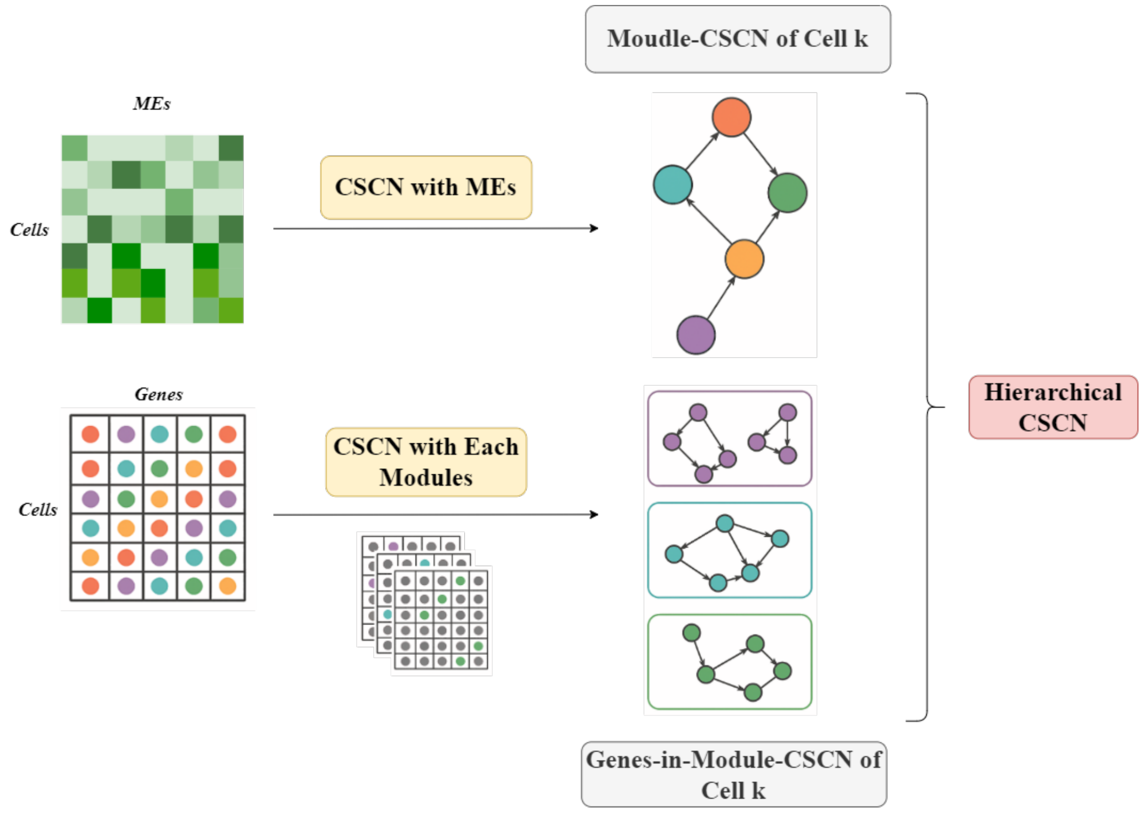


**Fig. S1.** Data preprocessing via WGCNA.

##### *S1.1.2. Dimension-reduced inference via NMF.*

When the objective is to transform noisy expression profiles into stable causal regulatory features rather than to recover explicit gene-level networks, we apply non-negative matrix factorization (NMF) to $X \in \mathbb{R}^{n\times m}$, $X \approx WH$ with $W \in \mathbb{R}^{n\times r}$, $H \in \mathbb{R}^{r\times m}$, $r \ll m$. Causal networks are constructed in the reduced module space of dimension $r$, yielding module-level CSCNs per cell; the resulting Causal Katz Matrix is then mapped back to gene resolution through the loading matrix $H$. The workflow is illustrated in Figure S2.

##### *S1.1.3. Targeted subset inference.*

For applications aimed at dissecting the regulatory relationships underlying specific biological processes—such as pathway-level causal intervention studies or estimation of process-specific treatment effects—we first curate a subset of genes or modules directly implicated in that process (e.g., based on pathway annotations, differential expression, or preliminary association screening) and

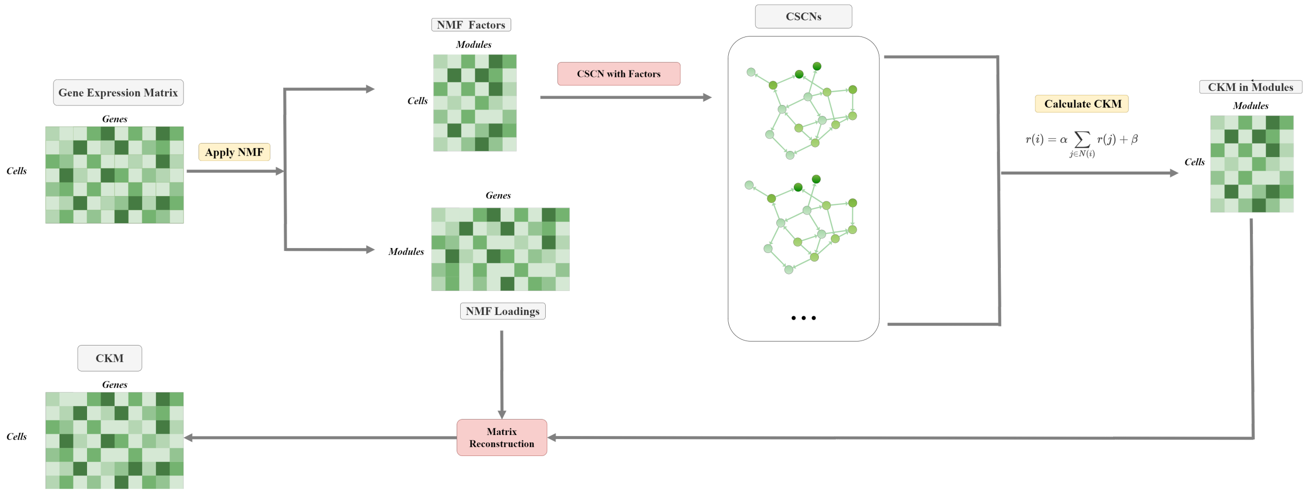


**Fig. S2.** Data preprocessing via NMF.

then confine the CSCN pipeline to this tailored set. This focused strategy both alleviates computational burden and filters out noise from unrelated components, thereby enabling more accurate causal recovery within the biological context of interest.

#### S1.2. Computational acceleration

The PC procedure is combinatorially and computationally prohibitive due to the exponential growth of conditioning sets and the cost of local neighborhood counts. To alleviate this, we first index all cell-wise expression profiles with a KD-tree, treating each cell as a point in the relevant expression subspace. Range-count queries for neighborhood construction leverage bounding-box pruning, reducing per-query cost from $O(nm)$ to approximately $O(n^{1-1/m})$ and thereby compressing the per-cell complexity to $O(m^2 2^m n^{1-1/m})$ in the full gene-level setting.

For further speedup, we layer on a bitmap-based intersection scheme with dynamic programming (DP) to efficiently compute conditional neighborhoods. Each gene (or module) and its local neighborhood is encoded as a bitset $B_g \in \{0, 1\}^n$, where bit $i$ is 1 if cell $i$ lies in the expression interval for gene $g$. The joint neighborhood for a conditioning set $S = \{z_1, \ldots, z_k\}$ is then the bitwise AND of its members:

$$B_S = \bigwedge_{z \in S} B_z.$$

We cache these intersections so that extensions of a known subset can be updated incrementally: for any $S$ and a new gene $z \notin S$,

$$B_{S \cup \{z\}} = B_S \ \& \ B_z,$$

requiring only one additional bitwise AND. This DP-style transition avoids recomputing full intersections from scratch when exploring overlapping conditioning subsets. Counting cells in the intersection is then a population count on $B_S$, and the cost of computing/intersecting $k$ neighborhoods reduces from $O(kn)$ to $O(k \cdot \lceil n/w \rceil)$, where $w$ is the machine word size (e.g., 64), since each bitwise operation processes $w$ bits simultaneously. Together, the KD-tree pruning and bitmap+DP intersection caching yield substantial practical runtime reductions, particularly under sparse conditioning regimes and overlapping subset exploration.

#### S1.3. Biomarker Discovery

Our discovery workflow (Fig. S3) first identifies candidate genes by applying DESeq2 for differential expression analysis on scRNA-seq data from normal and tumor cells. These candidates are then assessed within a Cell-Specific Causal Network (CSCN) framework. By introducing a virtual "Disease" node into the network, we calculate the Average Causal Effect (ACE) for each candidate gene's impact on the disease state. Only genes with a statistically non-zero ACE are validated as causal biomarkers, ensuring they have a direct influence on the phenotype beyond simple correlation.

#### S1.4. Downstream Analysis

Downstream Analysis (Fig. S4): Individual CSCNs are integrated to generate a global Causality Score Matrix (CKM) and global network for population-level insights. These integrated results facilitate functional subgroup identification, biomarker discovery, pseudotime reconstruction, and entropy analysis.

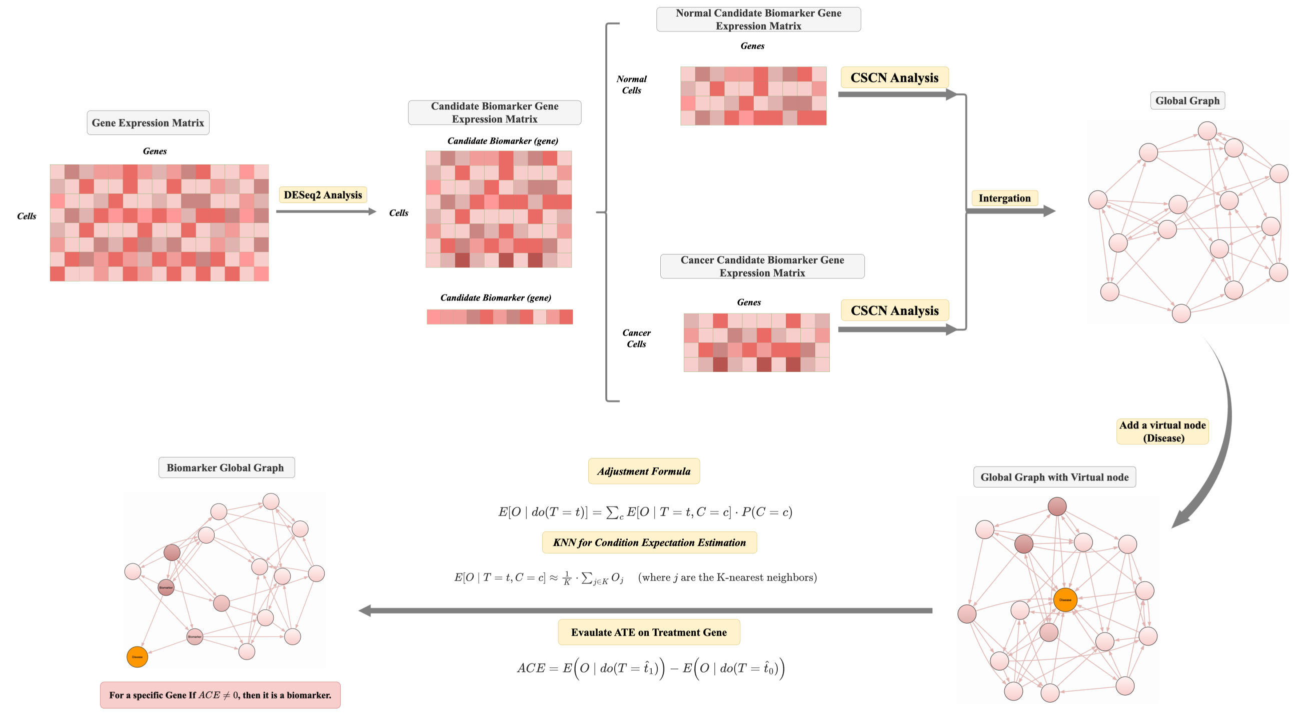


**Fig. S3.** Biomarker Discovery

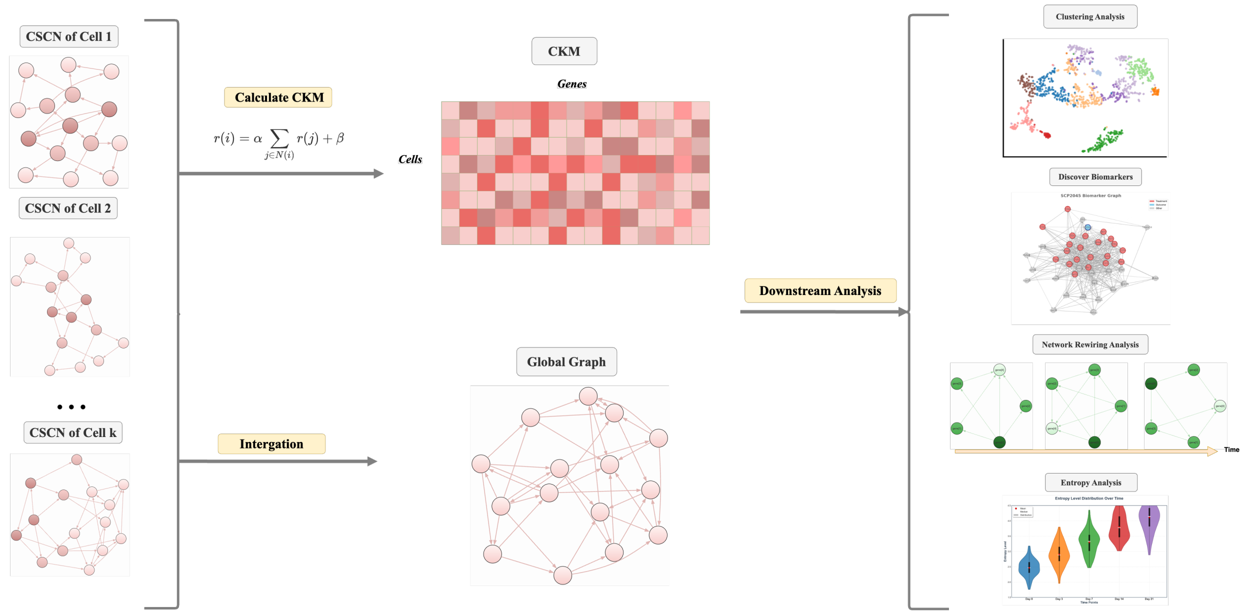


**Fig. S4.** Downstream Analysis of CSCN

**Table 1.** Adjusted Rand Index (ARI) obtained with different feature inputs

| **Method** | **Input** | E–GEOD 93593 | E–MTAB 8414 | E–MTAB 10434 | E–MTAB 9388 | E–GEOD 71585 | E–GEOD 75140 | E–MTAB 9067 | E–MTAB 5061 | E–ENAD 13 |
|---|---|---|---|---|---|---|---|---|---|---|
| KMeans | GEM | 0.16 | 0.60 | 0.01 | 0.13 | 0.26 | 0.27 | 0.08 | **0.40** | 0.13 |
| | NDM | 0.18 | 0.00 | -4.09 | 0.14 | 0.14 | 0.00 | 0.03 | 0.25 | 0.21 |
| | CNDM | 0.19 | 0.02 | 0.06 | 0.16 | 0.13 | **0.47** | 0.17 | 0.22 | **0.34** |
| | CKM | **0.27** | **0.68** | **0.09** | **0.42** | **0.30** | 0.40 | **0.25** | 0.31 | 0.32 |
| KMedoids | GEM | 0.17 | 0.23 | 0.01 | 0.15 | 0.29 | 0.05 | 0.05 | 0.21 | 0.07 |
| | NDM | 0.06 | 0.07 | 0.04 | 0.00 | 0.08 | 0.03 | 0.07 | **0.24** | 0.10 |
| | CNDM | 0.13 | 0.07 | 0.05 | 0.11 | 0.15 | 0.04 | 0.04 | 0.20 | 0.05 |
| | CKM | **0.26** | **0.58** | **0.71** | **0.24** | **0.29** | **0.33** | **0.17** | 0.22 | **0.27** |

**Note:** Boldface indicates the best result within each method–dataset pair.
